# Enzymatic stoichiometry is uncoupled with soil nutrient stoichiometry in highly-weathered Ferralsols across subtropical forests in China

**DOI:** 10.64898/2026.11.23.768509

**Authors:** Mingfeng Zhang, Andi Li, Xiangyin Ni

## Abstract

Ferralsols are highly weathered and nutrient limited, which may change microbial investment strategy for nutrient-acquiring enzymes for maintaining carbon (C) and nutrient stoichiometry. The equilibrium of soil C, nitrogen (N) and phosphorus (P) stoichiometry has well been identified across a broad range of terrestrial ecosystems, but it remains unclear whether enzymatic stoichiometry is coupled with soil nutrient stoichiometry in these highly-weathered Ferralsols with nutrient limitation, particularly in deep soils with less nutrient input. Here we collected topsoils (0-10 cm) and subsoils (40-50 cm) at 127 forest sites along 3000 km transects across subtropical Ferralsols in southern China to assess the enzymatic stoichiometry of β-1,4- glucosidase (BG), β-1,4-*N*-acetylglucosaminidase (NAG), leucine aminopeptidase (LAP) and acid phosphatase (AP). We found that soil BG, NAG, LAP and AP activities were significantly higher in topsoils than subsoils, and more energy was invested to N- and P-acquiring enzymes at both depths, suggesting that soil microbes were limited by both N and P in Ferralsols. Enzymatic stoichiometry was not significantly related to soil nutrient stoichiometry at both depths except for a positive correlation (*P*=0.029) between enzymatic and soil C/P ratios in subsoils. The nutrient-poor subsoils limit enzymatic production, resulting in soil microorganisms down-regulate enzymatic activity in these subtropical forests with fast C turnover. These results highlight that microbes may trade off the secretion of soil enzymes to acquire additional N and P in Ferralsols with high C (energy) investment in subtropical forests.

## Introduction

Soil enzymes are important participants in biogeochemical processes such as the decomposition of plant litter and soil organic matter (SOM), and changes in their activities are also directly related to the efficiency of microbial nutrient utilization (Deforest, 2009; Fanin *et al*., 2016). Soil microorganisms obtain nutrients from SOM by secreting specific enzymes, so that changes in soil enzymatic activities are generally used as important proxies to evaluate microbial nutrient requirements (Moorhead and Sinsabaugh, 2006). Therefore, profiling changes in soil enzymatic activities and stoichiometry can help to understand and predict nutrient turnover and SOM stabilization in response to resource constraints in soils (Cleveland and Liptzin, 2007; Kivlin and Treseder, 2013).

Soil enzymatic stoichiometry is the ratio of specific enzymatic activities associated with soil carbon (C), nitrogen (N) and phosphorus (P) acquisition, reflecting the resource requirements of microorganisms as well as the equilibrium of soil stoichiometry (Yang *et al*., 2020). Changes in soil enzymatic stoichiometry were regulated by microbial nutrient requirements, and the imbalanced response of soil nutrient supply and microbial growth requirements is usually characterized by high or low soil enzymatic production (Deng and Shangguan, 2017; He *et al*., 2020).

Microorganisms could secrete the appropriate nutrient-acquiring enzymes to mine nutrients occluded in SOM, and one enzyme can acquire several related nutrients (Mori, 2020). Sinsabaugh et al. (2008) found that the average enzymatic C:N:P ratios converges to 1:1:1 at a global scale, suggesting that soil enzymatic stoichiometry keep an equilibrium with soil C, N and P stoichiometry. Although this ratio is not fully consistent across differential ecosystems (Wang *et al*., 2014; Zheng *et al*., 2017), there are some studies found soil enzymatic stoichiometry is important in predicting soil nutrient limitation (Zhao *et al*., 2018; Bai *et al*., 2021).

Most studies only evaluated surface soil enzymatic stoichiometry above 0-15 cm depth, with few studies focus on subsoils (Eilers *et al*., 2012). Several studies have reported decreases in soil microbial community diversity, microbial biomass and soil enzymatic activity with increasing soil depth (Gelsomino and Azzellino, 2011; Herold *et al*., 2014). Topsoils are rich in organic matter with high microbial abundance, showing high levels of soil enzymatic activity (Lugosz and Piotrowska-Dlugosz, 2022). By contrast, the nutrient-poor subsoils limit microbial activity and the secretion of enzymes (Poffenbarger *et al*., 2020). In addition, differences in the degree of weathering and composition of parent minerals at different depths also affect soil nutrient content and microbial communities, which, in turn, alter soil enzymatic activity (Kögel-Knabner and Amelung, 2021; Marinari *et al*., 2021). For example, substrates adsorbed on mineral surfaces were not utilized quickly by microorganisms (Kaiser *et al*., 2007). At the same time, SOM could be bound with iron oxides, resulting in a decrease in nutrient availability (Scheel *et al*., 2007). These differential levels of nutrients in topsoils and subsoils could largely control enzymatic activity and stoichiometric equilibrium, but our current understanding is very limited.

Ferralsols in low-latitude areas are highly weathered, but the rich clay minerals and metal oxides adsorb a great proportion of nutrients. In soils rich in reactive iron oxides, organic P content decreases gradually with the degree of weathering (Fan *et al*., 2019; Basinski *et al*., 2024). Meanwhile, the binding of available nutrients to minerals further depletes soil dissolved N and P (Turner *et al*., 2014). Therefore, subtropical Ferralsols often show strong nutrient limitation, which could stimulate soil microorganisms to devote more resources to enzymatic production to regulate the imbalance between the limited soil resources and rapid metabolism requirements (Allison *et al*., 2007; Qiu *et al*., 2023).

Here we hypothesized that soil enzymatic C/N and C/P ratios are lower in topsoils than subsoils in highly-weathered Ferralsols and are uncoupled with soil C/N and C/P stoichiometry as high C investment. We measured soil C- (β-1,4- glucosidase, BG), N- (β-1,4-*N*-acetylglucosaminidase, BAG, and leucine aminopeptidase, LAP) and P- (acid phosphatase, AP) acquiring enzymatic activities and stoichiometry in topsoils (0-10 cm) and subsoils (40-50 cm) across 127 forest sites in Ferralsols in subtropical China.

## Materials and Methods

### Ferralsols forest transects

This study was conducted in two transects across subtropical forests in southern China (20-34°N, 110-120°E). The subtropical region covers about 25% of China’s land area. The natural vegetation is dominated by broad-leaved evergreen forests, and soils are classified as Ferralsols (China soil taxonomy) or Ultisols (USDA taxonomy). We selected the west-east (1000 km) and north-south (2000 km) transects of subtropical forests in southern China for field investigation (Fig. 1). The zonal broad-leaved evergreen forests with no or little human disturbance were used.

**Figure 1.**
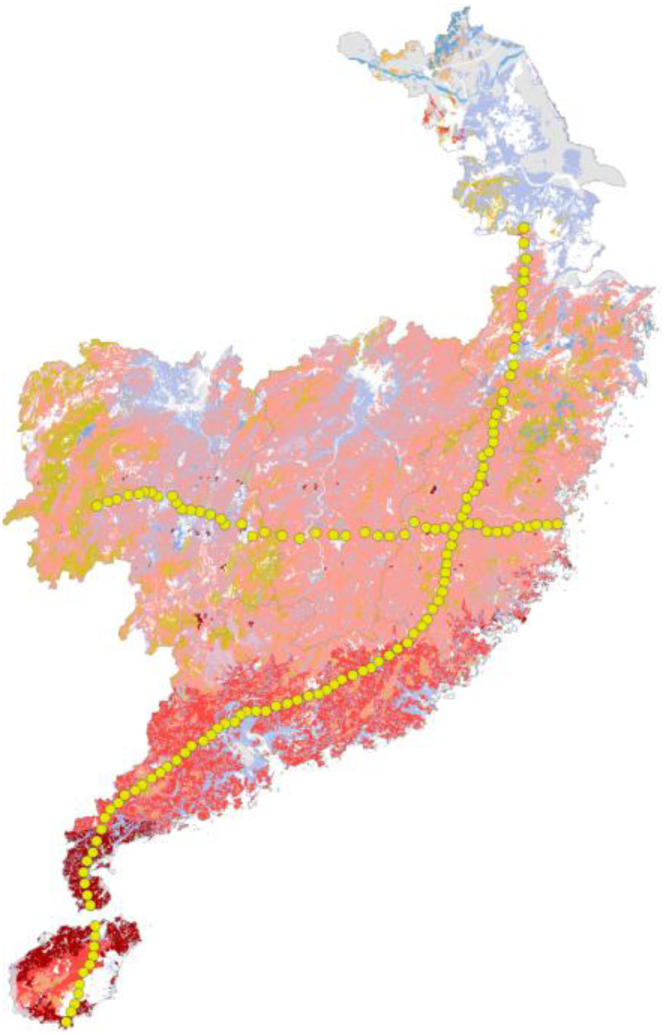
Field sampling sites in Ferralsols forest transects. The red areas show the distributions of Ferralsols in southern China, and the solid dots show sampling sites.

### Field sampling

Sampling plots of 20 m × 20 m in size were determined at each sampling site, and the longitude, latitude and elevation, as well as the parent material, soil types, forest types and dominant tree species were recorded. We selected the flat forest sites, removed the surface litter layer and collected topsoils (0-10 cm) and subsoils (40-50 cm) with mixtures from 3-5 soil profiles at 2 m away from the dominant trees. Soil samples were passed through a 2-mm sieve and stored at 4 °C for chemical analyses within two weeks.

### Soil enzymatic activities

Soil β-1,4-glucosidase (BG), β-1,4-*N*-acetylglucosaminidase (NAG) and acid phosphatase (AP) activities were determined by using *p*-nitrophenol as substrate (Kourtev, 2002; Dick, 2011). Briefly, 0.8 mL of acetate buffer solution (pH 5.5) and 0.2 mL of associated substrates were added to the soil homogenate, and the mixture was cultured at 37 °C for 3 h (acid phosphatase culture for 1 h). Then 0.2 mL 0.5 mol/L CaCl_2_ and 0.8 ml 0.5 mol/L NaOH solution were added, and the absorbance at 410 nm was measured. The soil leucine aminopeptidase (LAP) activity was determined by using L-leucine-*p*-nitroanilide as substrate (Sinsabaugh *et al*., 1999). Briefly, 60 μL L-leucine-nitroaniline solution was added, and the mixture was cultured at 30 °C for 1 h. Then, the solution was boiled for 5 minutes and centrifuged at 12000 rpm for 5 minutes before the absorbance of the supernatant was determined at 405 nm.

### Edaphic and climatic variables

Soil total carbon (TC), nitrogen (TN) and phosphorus (TP) concentrations were measured by the potassium dichromate-external heating method, the semi-micro Kjeldahl digestion method and the molybdenum anti-mony colorimetric method, respectively (Liu *et al*., 1996). Soil nitrate (NO_3_^−^) and ammonium (NH_4_^+^) concentrations were determined by extracting 5 g of fresh soils with 20 mL 2 mol/L KCl solution, and the extracts were measured by a Continuous Flow Analyzer (AA3, Bran + Luebbe, Germany; Ma *et al*., 2021). Soil dissolved organic carbon (DOC) and nitrogen (DON) were determined by a Shimadzu TOC-VcPh analyzer (Jones and Willett, 2006). Soil pH was measured by using a pH meter in a 1:2.5 soil: water (*v*/*v*) ratio (Men *et al*., 2023).

Soil microbial biomass C, N and P concentrations were measured with chloroform fumigation with conversion factors of 0.45 for C and 0.54 for N (Brookes *et al*., 1982; Brookes *et al*., 1985; Vance *et al*., 1987). Soil microbial biomarkers used in phospholipid fatty acids (PLFAs) are as follows: gram-positive bacteria (G^+^) (14:0 anteiso, 14:0 iso, 15:0 anteiso, 15:0 iso, 16:0 anteiso, 16:0 iso, 17:0 anteiso, 17:0 iso), gram-negative bacteria (G^−^) (15:1 ω5c, 16:1 ω7c, 17:1 ω8c, 18:1 ω5c, 18:1 ω7c, 18:1 ω9c), actinomycetes (ACT) (16:0 10-methyl, 17:0 10-methyl, 18:0 10-methyl), arbuscular mycorrhizal fungi (AMF) (16:1 ω5c) and fungi (18:2 ω6c) (Moore-Kucera and Dick, 2008; Dungait *et al*., 2011; Zheng *et al*., 2013; Orwin *et al*., 2018; Gorka *et al*., 2023). The mean annual temperature (MAT) and precipitation (MAP) were extracted from the WorldClim database based on the latitude and longitude of the sampling sites.

### Data analyses

Soil enzymatic stoichiometry was estimated according to the enzyme activities involved in C, N and P cycles.

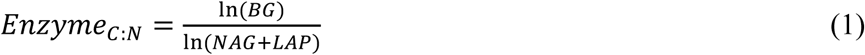

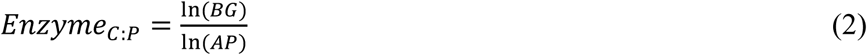

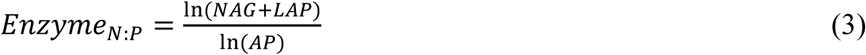

A random forest model was used to quantify the importance of climatic, edaphic and microbial properties on soil enzymatic stoichiometry. The variations of abiotic and biotic factors on enzymatic stoichiometry were determined by a redundancy analysis (RDA) by using the “*Vegan*” package in R. Correlations between soil enzymatic activities were analyzed with type Ⅱ standard major axis (SMA) regression by using the “*lmodel2*” package in R. Linear regression models were used to compare the relationship between enzymatic versus soil stoichiometry.

## Results

### Enzymatic activity and stoichiometry in Ferralsols

Soil BG, NAG, LAP and AP activities were higher at 0-10 cm than 40-50 cm depth (Fig. 2, *p*<0.05). At 0-10 cm depth, there were not significant correlations among the activities of the C-, N- and P-acquiring enzymes. The SMA regression slope associations between ln (BG) vs ln (NAG+LAP), ln (BG) vs ln (AP) and ln (NAG+LAP) vs ln (AP) were −1.12, 1.31 and 1.17, respectively (Fig. 3a-c). At 40-50 cm depth, there were positive correlations between ln (BG) and ln (NAG+LAP) and between ln (BG) and ln (AP) (Fig. 3a, b, *p*<0.05). The SMA regression slope associations between ln (BG) vs ln (NAG+LAP), ln (BG) vs ln (AP) and ln (NAG+LAP) vs ln (AP) were 0.97, 1.00 and 1.03, respectively (Fig. 3a-c).

**Figure 2.**
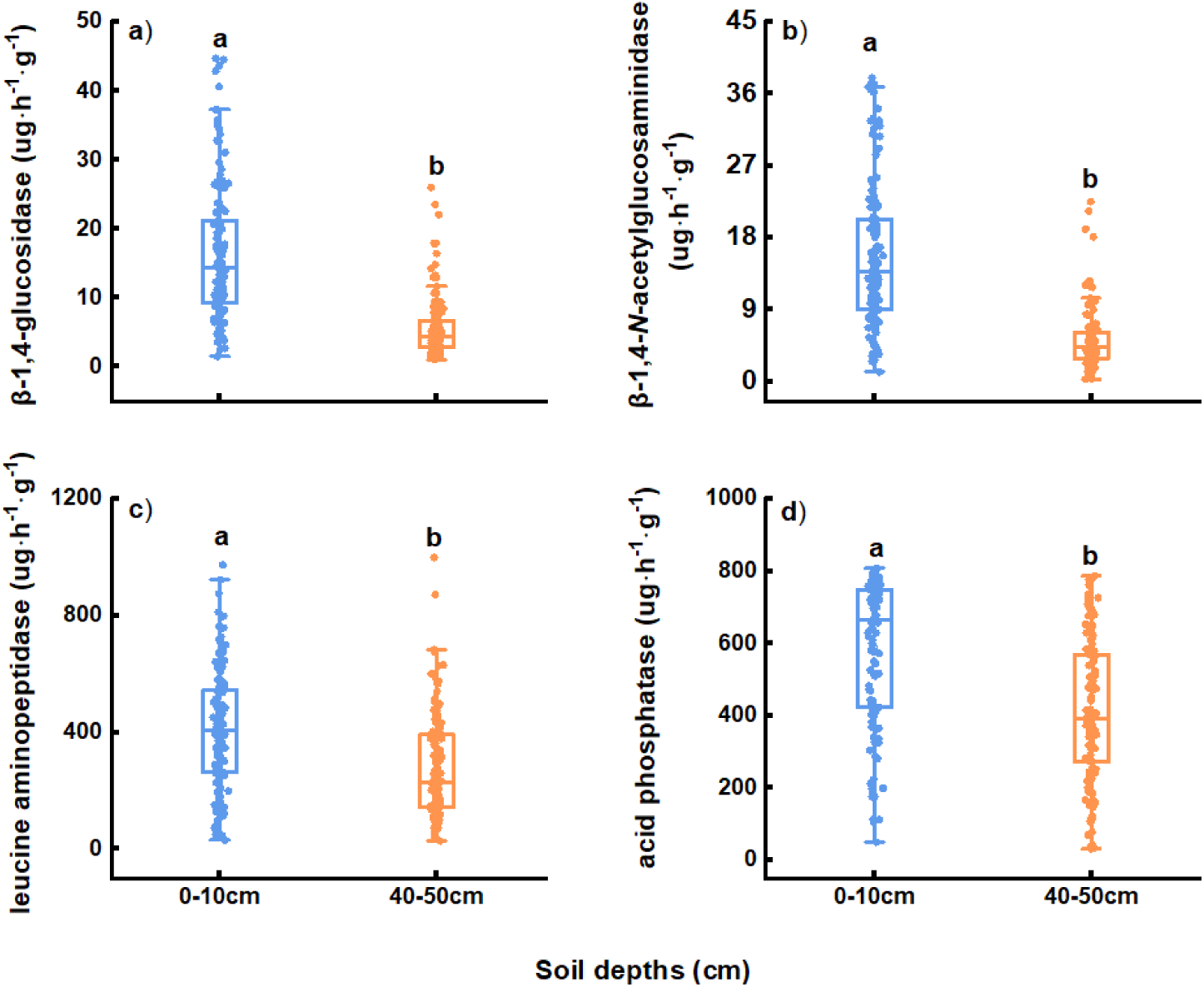
Changes in soil enzymatic activities at 0-10 cm and 40-50 cm depths. (**a**) β-1,4-glucosidase, (**b**) β-1,4-*N*-acetylglucosaminidase, (**c**) leucine aminopeptidase, (**d**) acid phosphatase. Different lowercase letters show significant differences in soil enzymatic activities between depths.

**Figure 3.**
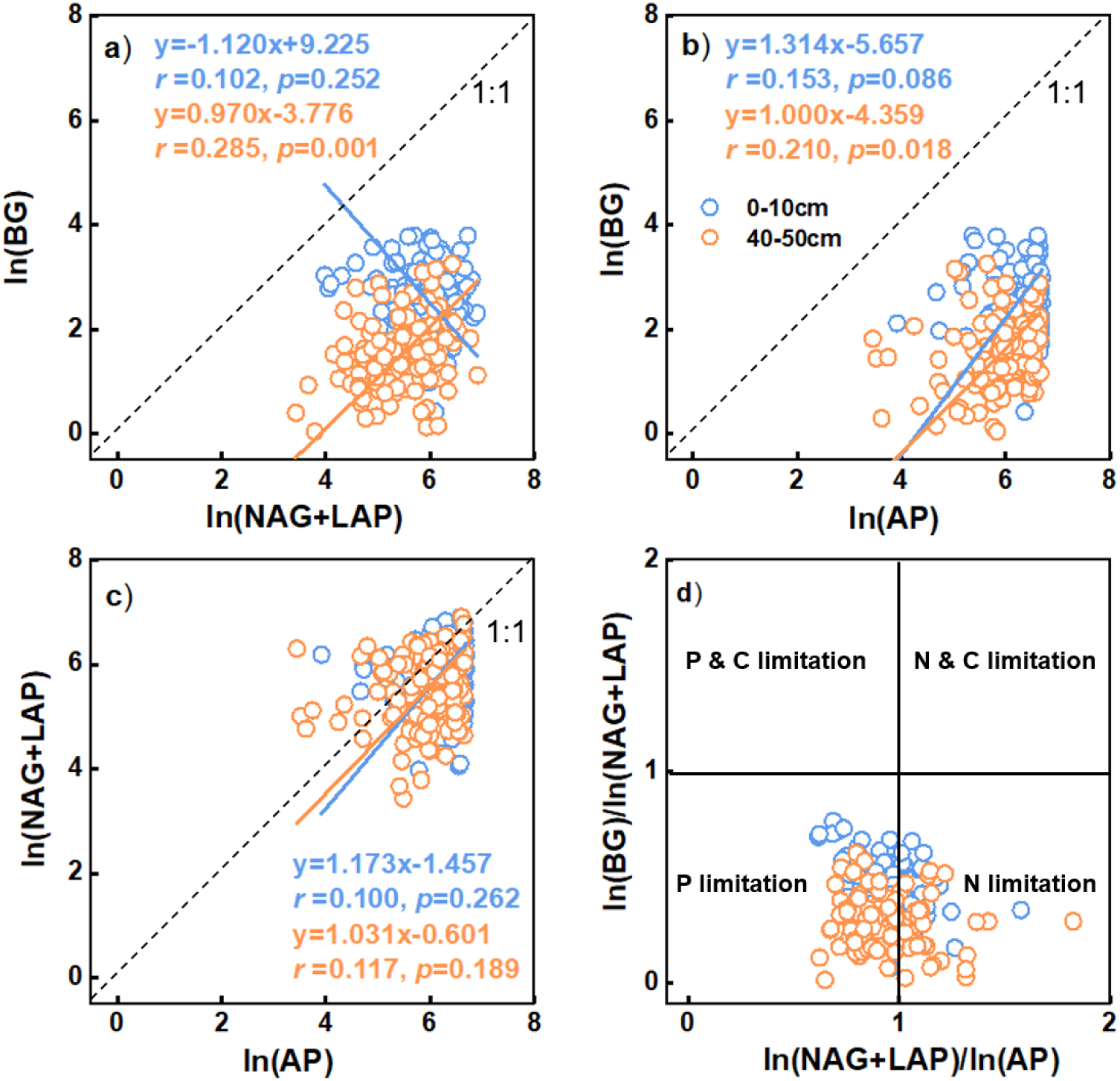
Standard major axis (type II) regression analysis between C-, N- and P- acquisition enzymatic activities at both soil depths. (**a**) ln (BG) vs ln (NAG+LAP), (**b**) ln (BG) vs ln (AP), (**c**) ln (NAG+LAP) vs ln (AP), (**d**) [ln (BG)/ln (NAG+LAP)] vs [ln (NAG+LAP)/ln (AP)]. The solid lines were the regression line. The dashed lines were the reference line with slope of 1. The regression equation, goodness of fit (*r*) and significance of models (*p*) were shown (*n*=127). \**p*<0.05, \*\**p*<0.01, \*\*\**p*<0.001.

However, in both depths, all SMA regression slopes were close to 1, suggesting that soil microorganisms invested more resources in N- and P-acquiring enzymes than in C-acquiring enzymes and that soil microbes were primarily limited by both N and P in subtropical forests (Fig. 3). Soil enzymatic C/N and N/P ratios were not correlated with soil C/N and N/P ratios (Fig. 4a, c), while soil enzymatic C/P ratio was positively correlated with soil C/P ratio (Fig. 4b, *p*<0.05), suggesting that soil AP activity decreased with decreasing soil P contents. This suggested that microbial secretion of N- and P-acquiring enzymes is a result of trade-off for practical resources rather than simple elemental access.

**Figure 4.**
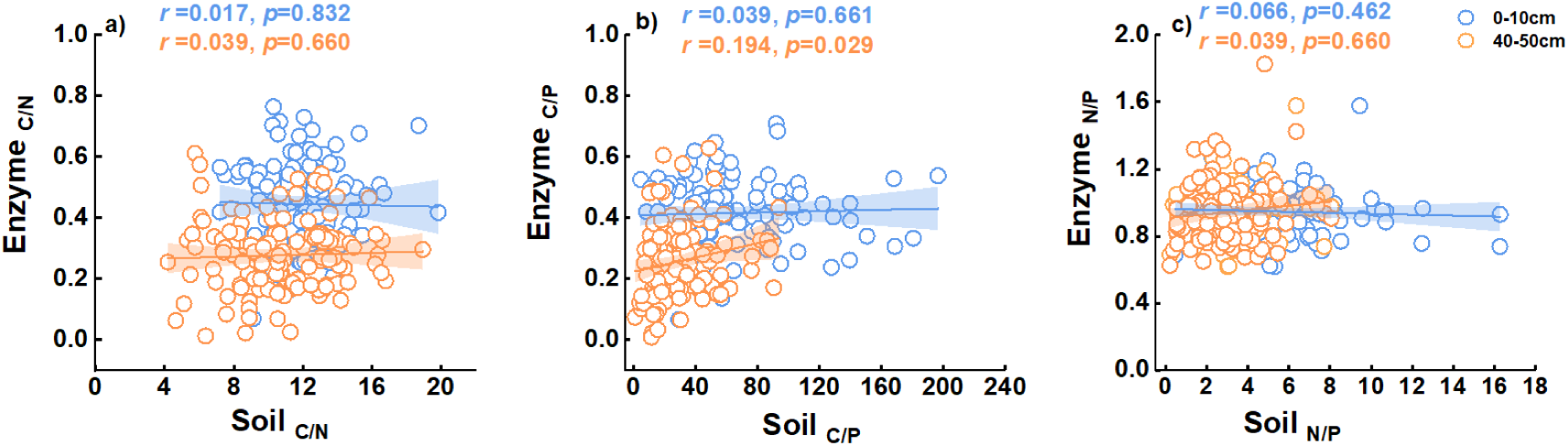
The linear relationship between enzymatic stoichiometry and soil nutrient stoichiometry at 0-10 and 40-50 cm soil depths. The fitting lines show slopes of linear regression model. Shaded bands indicated 95% confidence intervals (*n*=127). \**p*<0.05, \*\**p*<0.01, \*\*\**p*<0.001.

### Potential influences of soil enzymatic stoichiometry

In the 0-10 cm depth, the random forest analysis results showed that climatic, edaphic and microbial properties explained the variations in enzymatic C/N, C/P and N/P ratios by 26.42%, 26.18% and 10.03%, respectively (Fig. 5a-c). MAP was the most important variable in explaining soil enzymatic C/N and C/P ratios (Fig. 5a, b, *p*<0.01). In the 40-50 cm depth, climatic, edaphic and microbial properties explained the variations in enzymatic C/N, C/P and N/P ratios by 31.34%, 37.98% and 15.07%, respectively (Fig. 5d-f). The most important variables of soil enzymatic C/N and C/P ratios and enzymatic N/P ratio were MBP and G^−^ bacteria, respectively (Fig. 5a, b, *p*<0.01).

**Figure 5.**
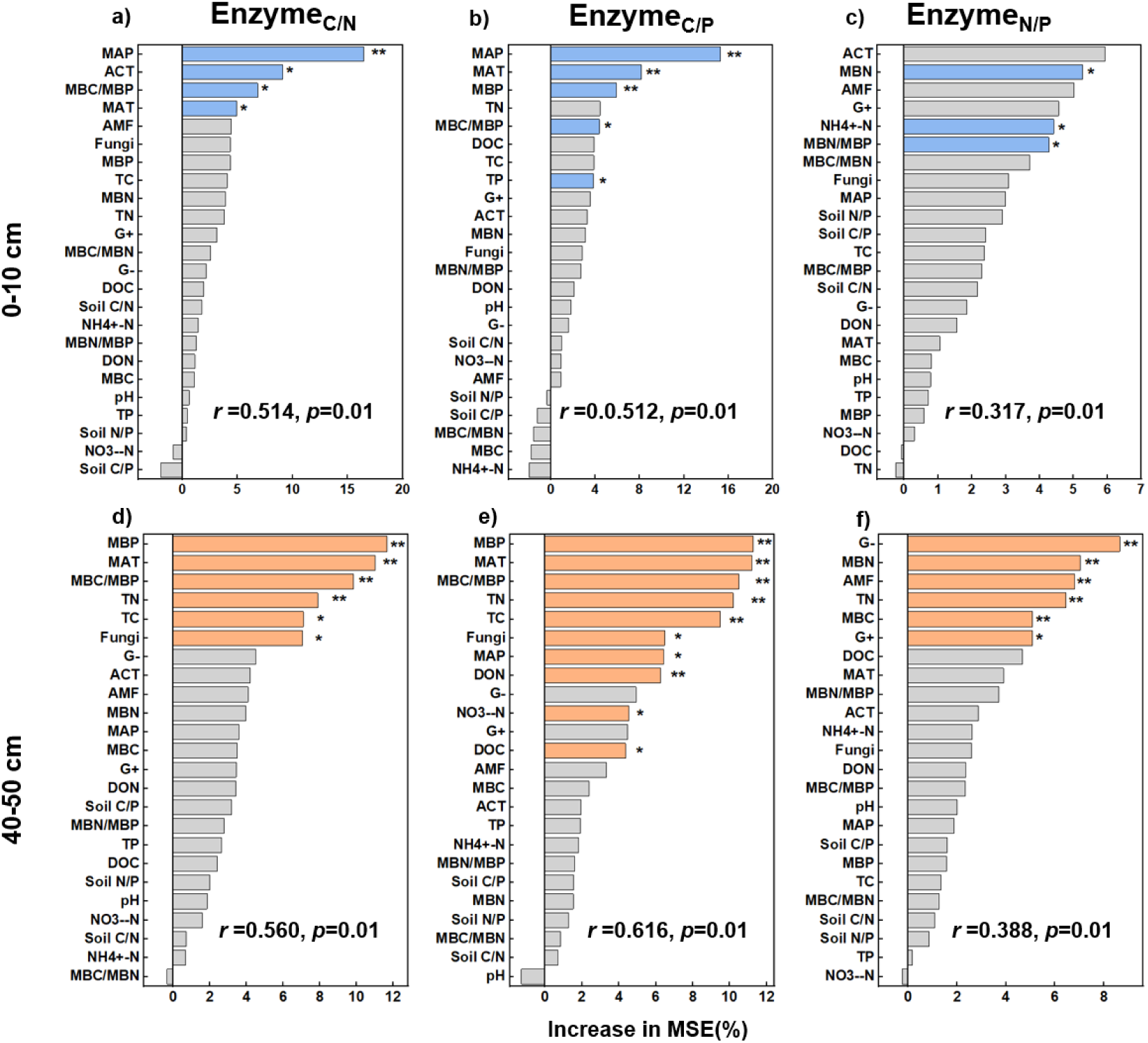
Random forest model shows importance of climatic, edaphic and microbial properties on soil enzymatic stoichiometry at (**a-c**) 0-10 and (**d-f**) 40-50 cm soil depths. Values represent the percentage increase in the mean square error (MSE). The goodness of fit (*r*) and significance of models (*p*) were shown. MAT: mean annual temperature, MAP: mean annual precipitation, NH_4_^+^-N: ammonium nitrogen, NO_3_^−^-N: nitrate nitrogen, TC: total carbon, TN: total nitrogen, TP: total phosphorus, DOC: dissolved organic carbon, DON: dissolved organic nitrogen, MBC: microbial biomass carbon, MBN: microbial biomass nitrogen, MBP: microbial biomass phosphorus, G^+^: gram-positive bacteria, G^−^: gram-negative bacteria, ACT: actinomycetes, AMF: arbuscular mycorrhizal. \**p*<0.05, \*\**p*<0.01, \*\*\**p*<0.001.

The redundancy analysis results showed that the total constrained variations in soil enzymatic stoichiometry at 0-10 and 40-50 cm depths were 39.84% and 29.05%, respectively (Fig. 6). Soil enzymatic stoichiometry was mainly associated with MAT, MAP, ACT, TC, NH_4_^+^, MBC/MBN and fungal PLFAs at 0-10 cm depth (Fig. 6a, *p*<0.05) and correlated with MAT, TC, DOC, MBP, ACT and G^+^ bacterial PLFAs at 40-50 cm depth across the Ferralsols transect (Fig. 6b, *p*<0.05). Particularly, MAP and MBP had the greatest influence on soil enzymatic stoichiometry at 0-10 and 40-50 cm depths, with explained variations of 13.45% and 10.32%, respectively.

**Figure 6.**
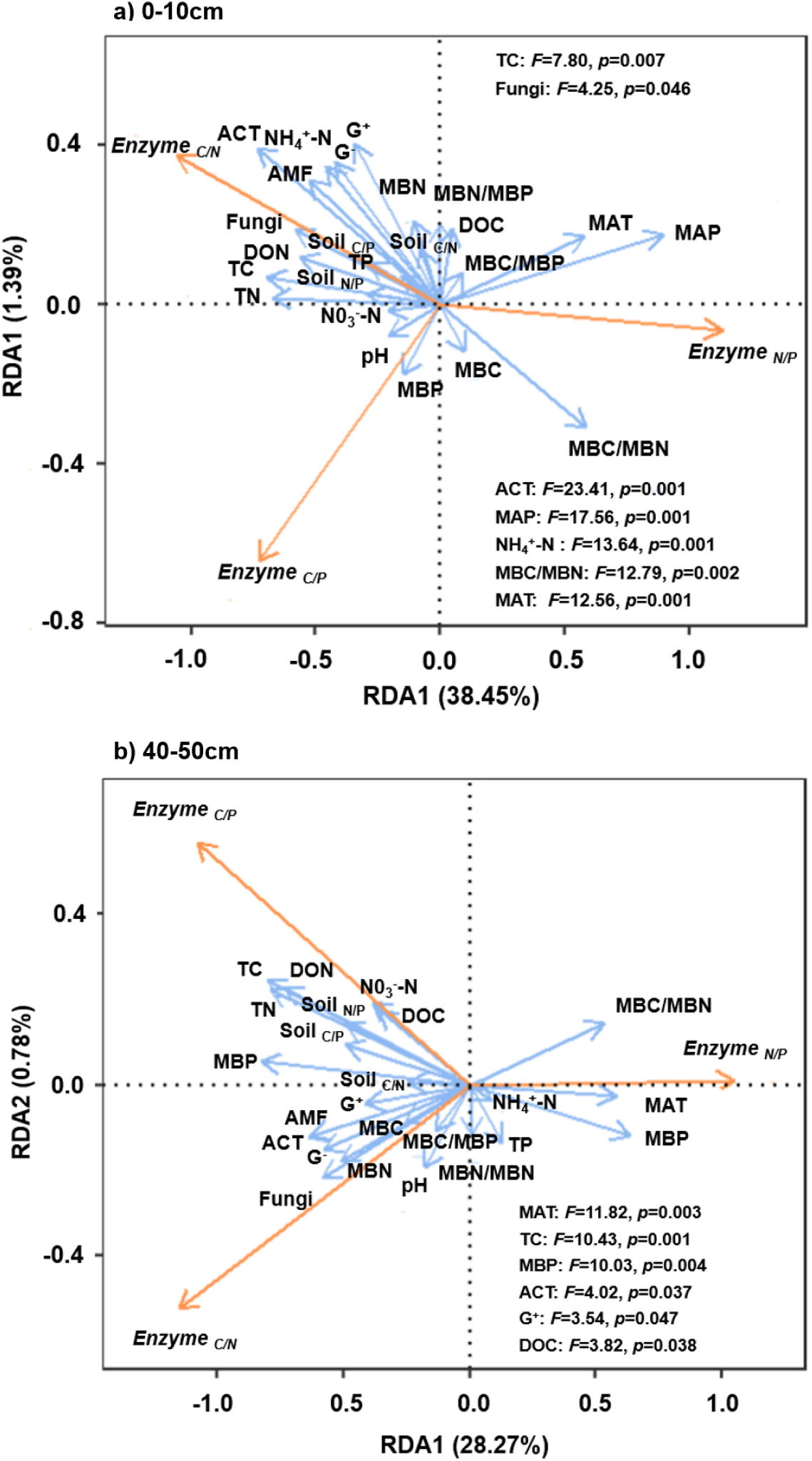
Redundancy analysis identifies the relationship among climatic, edaphic and microbial properties and soil enzymatic stoichiometry at (**a**) 0-10 and (**b**) 40-50 cm soil depths. Blue solid lines represent the environmental variables, and orange solid lines represent soil enzymatic stoichiometry (*n*=127). MAT: mean annual temperature, MAP: mean annual precipitation, NH_4_^+^-N: ammonium nitrogen, NO_3_^−^-N: nitrate nitrogen, TC: total carbon, TN: total nitrogen, TP: total phosphorus, DOC: dissolved organic carbon, DON: dissolved organic nitrogen, MBC: microbial biomass carbon, MBN: microbial biomass nitrogen, MBP: microbial biomass phosphorus, G^+^: gram-positive bacteria, G^−^: gram-negative bacteria, ACT: actinomycetes, AMF: arbuscular mycorrhizal.

## Discussion

Previous studies have found that soil enzymatic C/N ratio was lower in middle- and high-latitudes (Feng *et al*., 2019) because the increase in N reabsorption efficiency with continuous vegetation succession accelerates plants to take up more N from the soils, exacerbating soil N limitation, whereas P reabsorption efficiency was relatively stable (Zhang *et al*., 2022; Zhang *et al*., 2023). The highly-weathered Ferralsols in subtropical forests in low-latitudes have low soil P availability but high microbial demand for N to maintain resource equilibrium (Zhou *et al*., 2020). There were evidences that the differences in distribution of soil nutrients across soil depths might be related to a variety of environmental factors, such as atmospheric N deposition, climate variables, soil weathering and nutrient leaching (Markewitz *et al*., 2004; Phoenix *et al*., 2012; Chen *et al*., 2022).

Soil microorganisms exhibited co-limitation by both N and P in these highly-weathered Ferralsols (Fig. 3). This was likely ascribed to low P concentrations in long-term highly weathered subtropical forest soils as well as soil P adsorption by reactive minerals that reduce soil P availability, resulting in microorganisms showing a higher demand for soil P (Cleveland *et al*., 2002; Waring *et al*., 2014). Our results that MBP contributed the most to alter soil enzymatic C/N and C/P ratios at 40-50 cm depth also support this idea (Fig. 5d, e). Therefore, changes in soil MBP concentration may strongly stimulate soil microbes to devote available resources to P-acquiring enzymatic production to alleviate nutrient deficit, but this strategy also increases microbial demand for soil N (Xu *et al*., 2017; Feyissa *et al*., 2022). However, higher rates of N mineralization in subtropical forests also accelerate N loss (Zhou *et al*., 2003).

Interestingly, however, soil enzymatic stoichiometry was not significantly correlated with soil stoichiometry at both topsoils and subsoils, except for a positive correlation between soil enzymatic C/P ratio and soil C/P ratio at 40-50 cm depth (Fig. 4, *p*<0.05). On the one hand, our results suggested that the most important influences on the changes in soil enzymatic stoichiometry in subtropical forests were MAP and MBP (Figs. 5, 6, *p*<0.01). Therefore, changes in soil nutrient availability stimulate microbial secretion of extracellular enzymes to a lesser extent than MAP and MBP. On the other hand, several studies have reported a greater influence of soil parent material on enzymatic activities in subsoils as mineral differences could affect nutrient adsorption (Fontes and Weed, 1996; Blonska *et al*., 2016; Piotrowska-Dlugosz *et al*., 2022). Subtropical soils were rich in iron oxides that strongly adsorb available P, leading to the presence of strong P limitation in soils, particularly in highly-weathered Ferralsols (Fink *et al*., 2016; Xiong *et al*., 2022). Therefore, soil microorganisms might be more sensitive to changes in soil P availability than C content in these subtropical forests (Liu *et al*., 2019).

In consistent with the results reported in previous studies that soil enzymatic activity increases with decreasing soil nutrients, soil AP activity decreased with decreasing soil P content in this study (Fig. 4b). This was attributed to the fact that enzymatic synthesis was dependent on the consumption of large amounts of soil N (Chen *et al*., 2018). However, as the available nutrient contents decreased with increasing soil depth, the N content in subsoils is not sufficient to sustain enzymatic production (Cui *et al*., 2022). From a resource allocation perspective, resource inputs to P-acquiring enzymes are increased when relative P limitation occurs in soil microbes (Waring *et al*., 2014). However, in resource-poor subsoils, there was a risk that microorganisms would consume more resources than they acquire, which causes a change in microbial strategy of resource allocation and consequently a decline in P-acquiring enzymatic activity (Stone *et al*., 2014). Other studies have reported that decrease in phosphatase activity in subsoils is caused by the binding of enzymes to iron oxides, which alter the structural morphology and active sites of enzymes, ultimately decreasing the enzyme’s reaction (Stone and Plante, 2014; Turner *et al*., 2014).

## Conclusions

Our results found that C-, N- and P- acquiring enzymatic activities were higher in topsoils than in subsoils and that soil microbes were co-limited by both N and P in highly-weathered subtropical Ferralsols. More importantly, we present a new insight that there was no correlation between enzymatic stoichiometry and soil stoichiometry in subtropical Ferralsols. In summary, this study contributed to a new theoretical understanding of soil enzymatic stoichiometry with soil C, N and P stoichiometric equilibrium in highly-weather subtropical forests. Future studies could further explore the response of enzymatic stoichiometry to soil stoichiometry in whole-profiles of different ecosystems, particularly in nutrient-poor soils, to determine the nexus between soil enzymatic activity and nutrient availability.

## Acknowledgements

We appreciate all FFT participants for their contributions in field sampling, chemical analysis and data compilation.

## Data availability

The data that supports the findings of this study are available from the corresponding author on reasonable request.

